# The population history of northeastern Siberia since the Pleistocene

**DOI:** 10.1101/448829

**Authors:** Martin Sikora, Vladimir V. Pitulko, Vitor C. Sousa, Morten E. Allentoft, Lasse Vinner, Simon Rasmussen, Ashot Margaryan, Peter de Barros Damgaard, Constanza de la Fuente Castro, Gabriel Renaud, Melinda Yang, Qiaomei Fu, Isabelle Dupanloup, Konstantinos Giampoudakis, David Bravo Nogues, Carsten Rahbek, Guus Kroonen, Michäel Peyrot, Hugh McColl, Sergey V. Vasilyev, Elizaveta Veselovskaya, Margarita Gerasimova, Elena Y. Pavlova, Vyacheslav G. Chasnyk, Pavel A. Nikolskiy, Pavel S. Grebenyuk, Alexander Yu. Fedorchenko, Alexander I. Lebedintsev, Sergey B. Slobodin, Boris A. Malyarchuk, Rui Martiniano, Morten Meldgaard, Laura Arppe, Jukka U. Palo, Tarja Sundell, Kristiina Mannermaa, Mikko Putkonen, Verner Alexandersen, Charlotte Primeau, Ripan Mahli, Karl-Göran Sjögren, Kristian Kristiansen, Anna Wessman, Antti Sajantila, Marta Mirazon Lahr, Richard Durbin, Rasmus Nielsen, David J. Meltzer, Laurent Excoffier, Eske Willerslev

**Author notes:** These authors contributed equally to this work (Martin Sikora, Vladimir V. Pitulko, Vitor C. Sousa, Morten E. Allentoft).

## Abstract

Far northeastern Siberia has been occupied by humans for more than 40 thousand years. Yet, owing to a scarcity of early archaeological sites and human remains, its population history and relationship to ancient and modern populations across Eurasia and the Americas are poorly understood. Here, we analyze 34 ancient genome sequences, including two from fragmented milk teeth found at the ~31.6 thousand-year-old (kya) Yana RHS site, the earliest and northernmost Pleistocene human remains found. These genomes reveal complex patterns of past population admixture and replacement events throughout northeastern Siberia, with evidence for at least three large-scale human migrations into the region. The first inhabitants, a previously unknown population of “Ancient North Siberians” (ANS), represented by Yana RHS, diverged ~38 kya from Western Eurasians, soon after the latter split from East Asians. Between 20 and 11 kya, the ANS population was largely replaced by peoples with ancestry related to present-day East Asians, giving rise to ancestral Native Americans and “Ancient Paleosiberians” (AP), represented by a 9.8 kya skeleton from Kolyma River. AP are closely related to the Siberian ancestors of Native Americans, and ancestral to contemporary communities such as Koryaks and Itelmen. Paleoclimatic modelling shows evidence for a refuge during the last glacial maximum (LGM) in southeastern Beringia, suggesting Beringia as a possible location for the admixture forming both ancestral Native Americans and AP. Between 11 and 4 kya, AP were in turn largely replaced by another group of peoples with ancestry from East Asia, the “Neosiberians” from which many contemporary Siberians derive. We detect gene flow events in both directions across the Bering Strait during this time, influencing the genetic composition of Inuit, as well as Na Dene-speaking Northern Native Americans, whose Siberian-related ancestry components is closely related to AP. Our analyses reveal that the population history of northeastern Siberia was highly dynamic throughout the Late Pleistocene and Holocene. The pattern observed in northeastern Siberia, with earlier, once widespread populations being replaced by distinct peoples, seems to have taken place across northern Eurasia, as far west as Scandinavia.

---

Northeastern Siberia (the modern Russian Far East) is one of the most remote and extreme environments colonized by humans. Extending from the Taimyr Peninsula in the west to the Pacific Ocean in the east, and from the China/Russia border north to the Arctic Ocean, the region is home to dozens of diverse ethnolinguistic groups. Recent genetic studies of the indigenous peoples of this land have revealed complex patterns of admixture, which are argued to have occurred largely within the last 10 ky^1–3^. However, human populations have been in the region far longer^4,5^, raising the possibility of a deeper, yet unknown population history.

The earliest, most secure archaeological evidence of human occupation of the region comes from the artefact-rich, high-latitude (~70° N) Yana RHS site dated to ~31.6 kya (Figure 1), though there are recently-discovered traces of an even earlier human presence^4,6^. Yana RHS yielded a flake-based stone tool industry and highly developed bone and ivory artefacts, reminiscent of technologies seen in the Eurasian Upper Palaeolithic (UP), and southern Siberia (Extended Data Fig. 1)^5,7^. By the time of the Last Glacial Maximum (LGM) ~23-17.5 kya^8^, the coldest and driest stage of the late Pleistocene, the Yana people had disappeared. Assemblages from the LGM and especially later are dominated by a distinctive microblade stone tool technology, which spread in a time-transgressive manner north and east out of the Amur region^9,10^, but did not reach Chukotka or cross the Bering Land Bridge (Beringia) until the end of the Pleistocene, well after the initial peopling of the Americas. Changes in material culture continued into the Late Holocene, but it remains debated whether these successive archaeologically distinct cultural complexes represent in situ technological evolution, perhaps in response to changing glacial and post-glacial climates, or if these represent distinct groups of people. If the latter, it is unclear how the groups are related to each other, to contemporary Siberians, and to Native Americans, whose ancestors possibly emerged in this region, or at the very least traversed it en route to Beringia.

**Figure 1.**
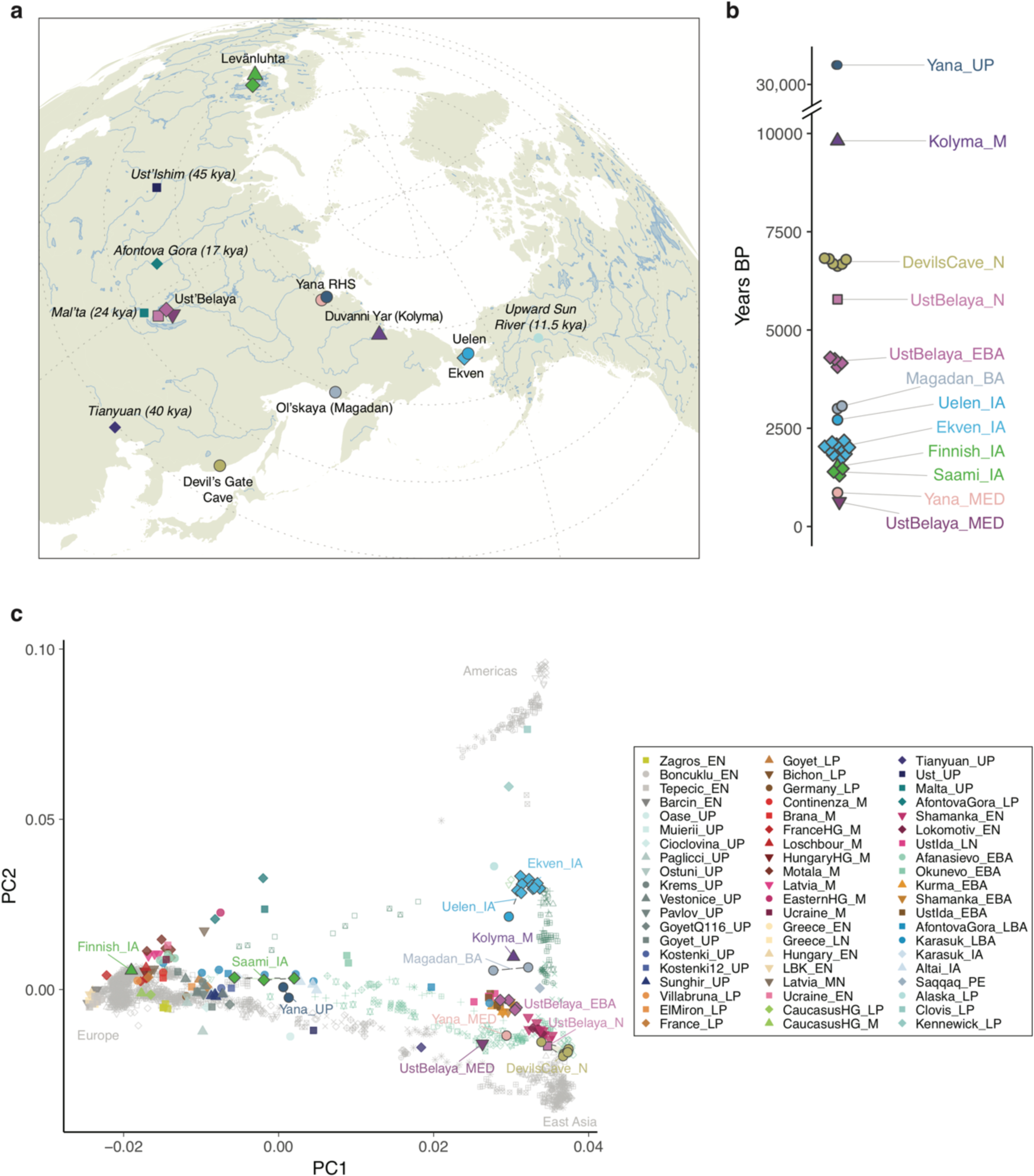
Genetic structure of ancient northeast Siberians. **a,** Sampling locations of newly reported and selected previously published individuals (italics). **b,** Sample ages. **c,** PCA of ancient individuals projected onto a set of modern Eurasian and American individuals. Abbreviations in group labels: UP – Upper Palaeolithic; LP – Late Palaeolithic; M – Mesolithic; EN – Early Neolithic; MN – Middle Neolithic; LN – Late Neolithic; EBA – Early Bronze Age; LBA - Late Bronze Age; IA – Iron Age; PE – Paleoeskimo; MED - Medieval

To investigate these questions, we generated whole-genome shotgun sequencing data from 34 ancient individuals, with radiocarbon ages ranging from 31,600 to 600 years before present (Figure 1; Extended Data Table 1; Supplementary Information 1-2; Supplementary Data Table 1). Our data include key samples from ancient individuals for the understanding of Siberian population history: two high-quality genomes (25X and ~7X coverage) sequenced from fragmented milk teeth (Supplementary Information 2) recovered from the Yana RHS site (Yana1 and Yana2, respectively), which are the oldest, northernmost Pleistocene human remains found to date; a high coverage genome (14X coverage) of an individual from the Duvanny Yar site near Kolyma River (Kolyma1), dated to 9.8 kya; fourteen genomes from ancient individuals from sites in far eastern Chukotka (Ekven, Uelen) and the northern coast of the Sea of Okhotsk (Ol’skaya), ranging from 3 kya to 2 kya; six individuals from the ~7.6 kya site of Devil’s Gate Cave in Primorskoye, northern East Asia^11^; as well as ten genomes from later, Holocene-age individuals from northern and southern Siberia (Ust’Belaya, in the Lake Baikal region, and Young Yana, which is from the same area but a different locality than Yana RHS), and from northwestern Eurasia (Finnish, Saami). We merged these data with large panels of previously published ancient and modern individuals (Supplementary Information 3; Supplementary Data Table 2)^3,12–42^.

## Ancient North Siberians from the Upper Palaeolithic Yana RHS

The Yana RHS human remains represent the earliest direct evidence of human presence in northeastern Siberia, a population we refer to as “Ancient North Siberians” (ANS). Both Yana RHS individuals were unrelated males, and belong to mitochondrial haplogroup U, predominant among ancient West Eurasian hunter-gatherers, and to Y chromosome haplogroup P1, ancestral to haplogroups Q and R, which are widespread among present-day Eurasians and Native Americans^43,44^ (Extended Data Fig. 2; Extended Data Table 1; Supplementary Information 4, 5). Genetic clustering of the individuals using outgroup-*f*_3_ statistics demonstrates broad genetic similarities with a wide range of present-day populations across Northern Eurasia and the Americas. This contrasts with other UP Eurasians such as those from Sunghir and Tianyuan, who share overall similar amounts of genetic drift with present-day populations but were geographically more restricted to West Eurasia and East Asia, respectively (Extended Data Fig. 3). Symmetry tests using *f*_4_ statistics reject tree-like clade relationships with both Early West Eurasians (EWE; Sunghir) and Early East Asians (EEA; Tianyuan); however, Yana is genetically closer to EWE, despite its geographic location in northeastern Siberia (Extended Data Fig. 3d, Extended Data Table 2; Supplementary Information 6). Using admixture graphs (*qpGraph*) and outgroup-based estimation of mixture proportions (*qpAdm*), we find that Yana can be modelled as EWE with ~25% contribution from EEA (Figure 2; Extended Data Fig. 3; Supplementary Information 6). Further estimates from demographic modelling of the high-coverage individual Yana1 using a site frequency spectrum (SFS)–based framework indicate an early divergence and mixture of the Yana lineage at ~39 kya (32.2-45.8), receiving 29.2% (21.3-40.1) contribution from EEA, which likely occurred very soon after the divergence of West Eurasians (EWE) and East Asians (EEA) (33.4-48.6 kya) (Figure 3a; Supplementary Information 7). Thus, Yana represents a distinct lineage among early Eurasians with affinities to both EWE and EEA, potentially a consequence of ancestral population structure, as also suggested by the presence of EEA ancestry and mitochondrial haplogroup M in early West Eurasia at Goyet 35 kya. We also estimate ~2% of Neanderthal ancestry in Yana, which is contained in longer genomic tracts than that of present-day individuals, similar to other UP Eurasians and consistent with the age of the Yana RHS remains (Supplementary Information 6)^32,39^.

**Figure 2.**
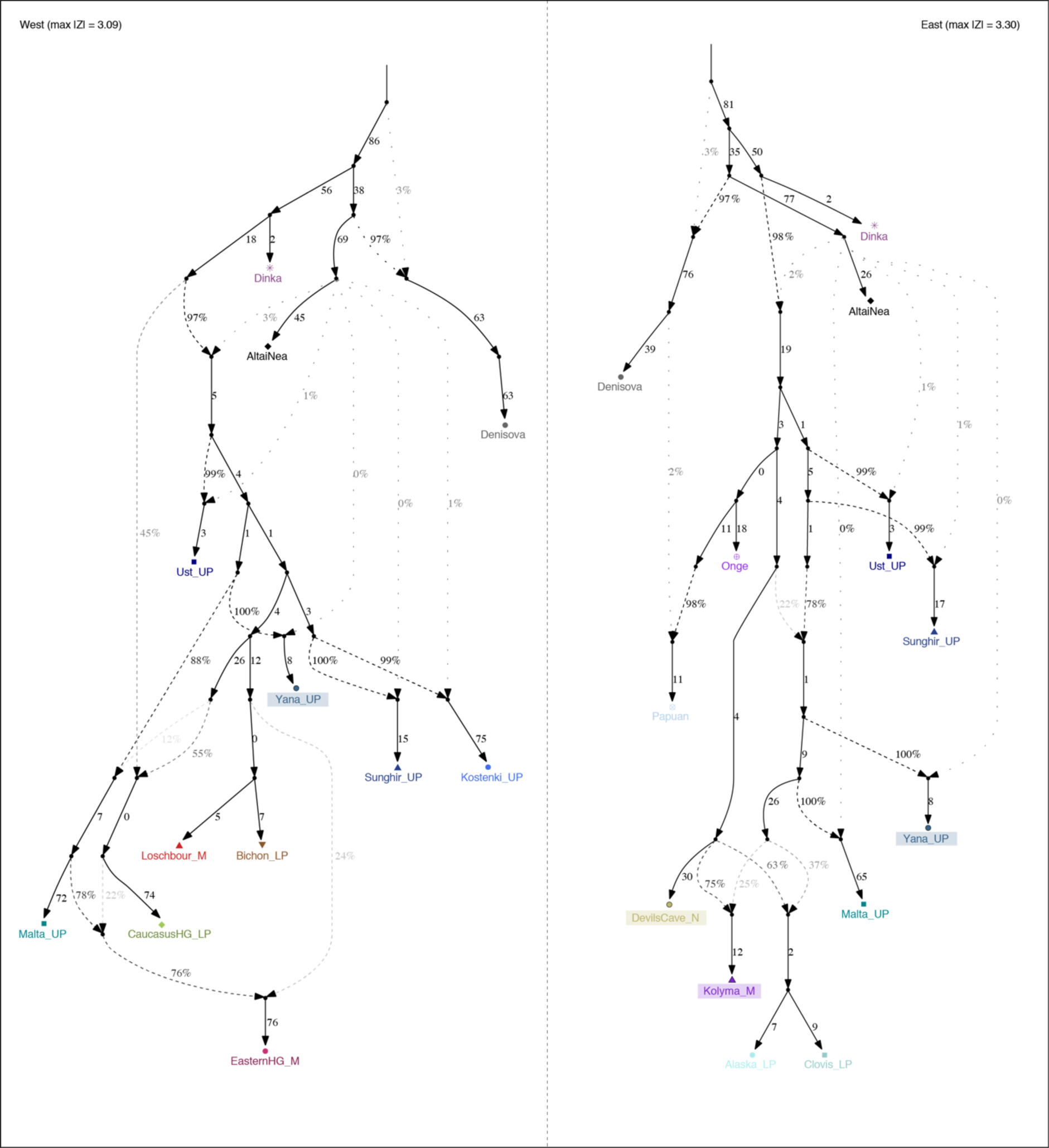
Admixture graph models of ancient and modern populations for western Eurasia (left) and East Asia and the Americas (right). Newly reported individuals are highlighted with coloured background. Early Upper Palaeolithic individuals were modelled allowing for a possible additional Neanderthal contribution to account for higher level of Neanderthal ancestry (dotted lines).

We next investigated how Yana relates to another enigmatic ancient Siberian population, the “Ancestral North Eurasians” (ANE) represented by the 24 kya Mal’ta individual from the Lake Baikal region, from which Native Americans were shown to derive ~40% of their ancestry^15^. Among all ancient individuals, Yana shares the most genetic drift with Mal’ta, and f4 statistics show that Mal’ta shares more alleles with Yana than with EWE (e.g. f4(Mbuti, Mal’ta;Sunghir, Yana) = 0.0019, Z = 3.99; Extended Data Table 2; Supplementary Information 6). Mal’ta and Yana also exhibit a similar pattern of genetic affinities to both EWE and EEA, consistent with previous studies^40,46^ (Extended Data Fig. 3e). The ANE lineage can thus be considered a descendant of the ANS lineage, demonstrating that by 31.6 kya early representatives of this lineage were widespread across northern Eurasia, including far northeastern Siberia. Fitting Yana and Mal’ta in an extended admixture graph including other EWE individuals reveals an increase in complexity of the ANS lineage over time due to differentiation and admixture with other lineages, with evidence for low levels of early Caucasus hunter-gatherer (CHG)–related ancestry in Mal’ta, as well as Western hunter-gatherer (WHG) and CHG–related ancestry in the individual from Karelia, commonly referred to as “Eastern hunter-gatherer” (EHG)^47^ (Figure 2, left).

The Yana high coverage genomes of two contemporaneous individuals provide an opportunity to investigate relatedness and levels of inbreeding at this remote UP settlement. We find that the two were not closely related, despite their similar mtDNA and Y chromosome haplogroups (Extended Data Fig. 4). Furthermore, analysing runs of homozygosity in the two individuals reveals no signatures of recent inbreeding in their genomes. Using data obtained from coalescent simulations, we estimate that the population at Yana is consistent with a recent effective population size of up to 500 individuals, remarkably high considering the extreme remoteness of the site. Our results mirror those observed at Sunghir, an early (~34 kya) European UP site located ~4,500 km southwest of Yana, reinforcing the view that wide-ranging networks of mate exchange and gene flow were present among UP foragers on the pre-LGM landscape^39^.

## Ancient Paleosiberians and the origins of Native Americans

Following the occupation at Yana RHS, there is an absence of archaeological sites in northeastern Siberia that lasts until the end of the LGM, when groups bearing a very distinctive stone tool technology appear in the region (~18 kya). It was within that intervening period that the ancestral Native American population emerged^15,41^, but prior to this study no genomes from individuals dating from this time period had been recovered in northeastern Siberia. We resolve this absence with the 9.8 kya Kolyma1 individual, representing a group we term “Ancient Paleosiberians” (AP). Our results indicate that AP are derived from a first major genetic shift observed in the region (Extended Data Fig. 5). Principal component analysis (PCA), outgroup *f*_3_-statistics and mtDNA and Y chromosome haplogroups (G1b and Q1a1a, respectively) demonstrate a close affinity between AP and present-day Koryaks, Itelmen and Chukchis, as well as with Native Americans (Extended Data Fig. 5; Supplementary Information 6). While these populations predominantly derive their ancestry from East Asia, admixture modelling shows that Kolyma1 derives from a mixture of EEA and ANS ancestry similar to that found in Native Americans, although Kolyma1 carries a greater EEA contribution (75% versus 63%) (Figure 2; Supplementary Information 6). For both AP and Native Americans, ANS ancestry appears more closely related to Mal’ta than Yana, therefore rejecting a direct contribution of Yana to later AP or Native American groups.

We then estimated the demographic parameters of population history models including Kolyma1 as well as the recently identified Ancient Beringians^41^ (Upward Sun River 1 [USR1]) and present-day Native Americans (Karitiana). We find that the ancestors of Kolyma1, Ancient Beringians and Native Americans diverged ~30 kya (26.8-36.4) from present-day East Asians (Han), in agreement with previous results^41^, with a subsequent divergence of Kolyma1 from the Ancient Beringian / Native American ancestral population at ~24 kya (20.9-27.9) (Figure 3; Supplementary Information 7). Both Kolyma1 and Native American ancestors received ANS-related gene flow at a similar time (Kolyma1 20.2 kya (15.5-23.7); USR1 19.7 kya (13.3-23.5)), which we model as a contribution from a population related to Yana. This gene flow amounts to 16.6% (7.5-22.2) of Yana-related ancestry into Kolyma1, and 18.3% (9.8-20.3) into USR1, comparable to the estimates obtained from the admixture graph modelling. We also tested an alternative model with a single admixture pulse in the ancestral population of Kolyma1 and USR1. While that model showed a comparable likelihood (Supplementary Information 7), results from the admixture analyses suggest differences in the ANS ancestry proportions between Kolyma1 and USR1, thereby favouring the two-independent pulses model. Kolyma1 thus represents the closest relative to the ancestral Native American population in northeastern Siberia known to date.

**Figure 3.**
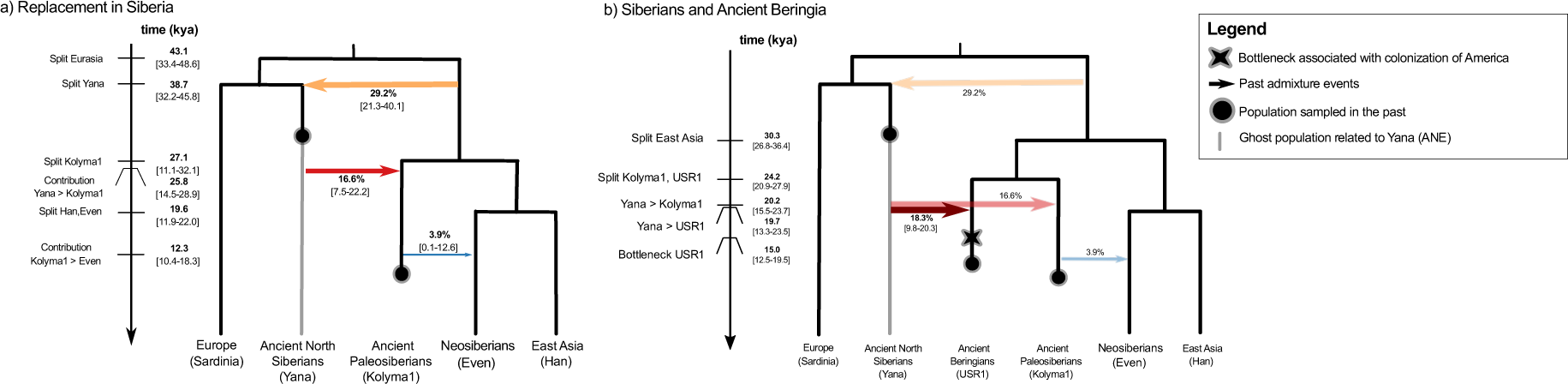
Demographic modelling of Siberian and Native American populations. Inferred parameters for models with: **a,** Ancient and modern Siberian populations, **b,** Siberian and Ancient Beringian. To account for confounding factors affecting estimates of admixture contributions we modelled explicitly: different sampling times for Yana and Kolyma1, different effective sizes for each population, potential bottlenecks representing founder events associated with each population split events, and in the ancestor of all Eurasians, a period of continuous migration between Europe and East Asian, and differential Neanderthal contribution into Europe and East Asia. The Neanderthal contribution was modelled by considering an unsampled (“ghost”) Neanderthal population contributing 3% into the ancestors of all Eurasian populations, and an extra 0.5% into the Asian lineage. Neanderthal effective size and split times were fixed according to recent estimates based on genome-wide SFS^79^. Shaded arrows for the “Siberia and Acient Beringia” model (**b**) indicate admixture proportions that were fixed to values estimated under model (**a**).

Changes in climatic conditions are commonly put forward as a principal driver of Pleistocene population movement and regional abandonment in Siberia. We used paleoclimatic modelling to infer geographic locations suitable for human occupation during the Pleistocene to further investigate this hypothesis. Prior to the LGM, when humans were present at Yana RHS, we infer that climatic conditions were suitable for human occupation across a large stretch of the Arctic coast of northeastern Siberia (Extended Data Fig. 8; Supplementary Information 8). Conditions in the region became less suitable during the LGM, supporting the lack of genetic continuity between Yana RHS and later groups. Interestingly, we find evidence for a potential refuge in southeastern Beringia, as well as a possible coastal corridor along the Sea of Okhotsk and the Kamchatka Peninsula during the LGM (e.g. panel 22ka, Alaska, Extended Data Fig. 8a), in line with previous reports^48^. A possible scenario for the ANS gene flow during the formation of Native Americans and AP might therefore have involved early ANS surviving in southeastern Beringia during the LGM, with subsequent admixture with EEA arriving by a coastal migration ~20 kya. This scenario would also be consistent with a divergence of Ancient Beringians from ancestral Native Americans in eastern Beringia rather than in Siberia, which is also supported by genetic data (Scenario 2 in^41^). Alternatively, the closer affinity of both Kolyma1 and Native Americans to Mal’ta rather than Yana could suggest a more southern location (Lake Baikal region) for the admixture, with a subsequent northward expansion following the LGM. While the archaeological evidence of a movement south during the LGM supports this scenario, the genetic isolation between Asians and ancestral Native Americans after ~24 kya would have been difficult to maintain if several populations sought refuge in the region. Our results nevertheless support the broader implication that glacial and post-glacial climate change was a major driver of human population history across Northern Eurasia.

## Holocene transformations across Siberia and Beringia

Recent genetic studies have documented additional, Holocene-age migrations and gene flow across the (now) Bering Strait involving both Paleoeskimos and Neoeskimos. Our genomic data provide further insights into the peoples involved, as well as the timing of these events. The 4 kya Saqqaq individual from Greenland^12^, representing Paleoeskimos, clusters with Kolyma1, but shows greater affinity to East Asians (Figure 1; Extended Data Table 2). Modelling Saqqaq as a mixture of Kolyma1 and Devil’s Gate Cave (DGC), we estimate it harbours around 20% DGC-related ancestry (Extended Data Fig. 7; Supplementary Information 6; Supplementary Data Table 3). Individuals from the Uelen and Ekven Neo-Eskimo sites (2.7 – 1.6 kya), located on the Siberian shore of the Bering Sea, closely resemble contemporary Inuit (Figure 1, Extended Data Fig. 6). We successfully fit them as a mixture of 69% AP (Kolyma1) and 31% Native American (Clovis) ancestry, thereby documenting a ‘reverse’ gene flow across the Bering Sea from northwestern North America to northeastern Siberia, in accordance with the linguistic back-migration of Eskimo-Aleut (Extended Data Table 2; Supplementary Information 6, 9; Supplementary Data Table 1). The source population of this gene flow post-dates the divergence of USR1 from other Native Americans (~20.9 kya^41^), as the Iron Age individuals at Ekven share more alleles with ancient Native Americans (Anzick-1, Kennewick) than with ancient Beringians (USR1), confirming previous results from present-day Inuit^49^ (Extended Data Table 2). To investigate the time of gene flow we performed linkage-disequilibrium (LD) based admixture dating^50^, using Saqqaq and Anzick-1 as source populations. We find a significant weighted LD curve with an estimated admixture date between 100 – 200 generations ago, depending on the data set (Supplementary Information 6). While these estimates show a considerable margin of error due to small sample size and limited amount of data for the Ekven population, they nevertheless suggest that gene flow related to Native Americans occurred back from the Americas into Siberia well after the disappearance of Beringia, but possibly as early as ~5 kya (~ 100 generations prior to the age of the earliest individual from Uelen and Ekven). That process contributed to a major ancestry component of contemporary Inuit populations. A genetic link has also been observed between North American populations speaking Na-Dene languages (Athabascans) and Siberian populations^51^. It has been suggested that this link reflects gene flow from a Paleoeskimo source represented by Saqqaq^52^, but a more recent study found evidence for a ghost source population more closely related to Koryaks^41^. Both admixture graph modelling (Supplementary Information 6) and chromosome-painting symmetry tests (Extended Data Fig. 5) show that Kolyma1 is a better proxy for this ghost ancestry than Saqqaq, therefore providing additional evidence against a contribution via a more recent migration of Paleoeskimos.

The Holocene archaeological record of northeast Siberia is marked by further changes in material culture. We used a temporal transect of ancient Siberians from ~6 kya to 500 years ago to investigate whether these cultural transitions were associated with genetic changes. We find that in a PCA of modern non-African populations, most contemporary Siberian populations are arranged along two separate genetic clines. The majority of individuals (here referred to as “Neosiberians”) fall on an East-West cline stretched out along PC1 between European populations at one end, and East Asian individuals including the ancient individuals from Devil’s Gate Cave at the other (Figure 1). A secondary cline between East Asians and Native Americans along PC2 includes Paleosiberian speakers and Inuit populations. Although AP ancestry (Kolyma1) was still common in other Siberian regions during the early Bronze Age (Extended Data Fig. 7), by the late Bronze Age we find it was largely restricted to the northeast, exemplified by a 3 kya individual from Ol’skaya (Magadan) that closely resembles present-day Koryaks and Itelmens. Using modern Even individuals to represent Neosiberians in our demographic model, we find evidence for a recent divergence from East Asians ~13 kya, with only low levels (~6%) of AP gene flow at ~11 kya (Figure 3; Supplementary Information 7). Thus, our data provides evidence for a second major population turnover in northeastern Siberia, with Neosiberians arriving from the south largely replacing AP, a pattern that is also evident in chromosome painting analyses of modern populations (Figure 4). Notable exceptions are populations such as the Ket, an isolated population that speaks a Yeniseian language and which has previously been described as rich in ANE-ancestry and with genetic links to Paleoeskimos^51^. The Ket fall on a secondary cline parallel to Neosiberians in the chromosome painting analysis and carry ~40% of AP ancestry (Figure 4; Extended Data Fig. 6). Our findings therefore reconcile the proposed linguistic link between the Yeniseian speaking Ket and Na-Dene speaking Athabascan populations (Supplementary Information 9), through shared ancestry with an AP metapopulation that at one time was more widespread across Northern Eurasia.

**Figure 4.**
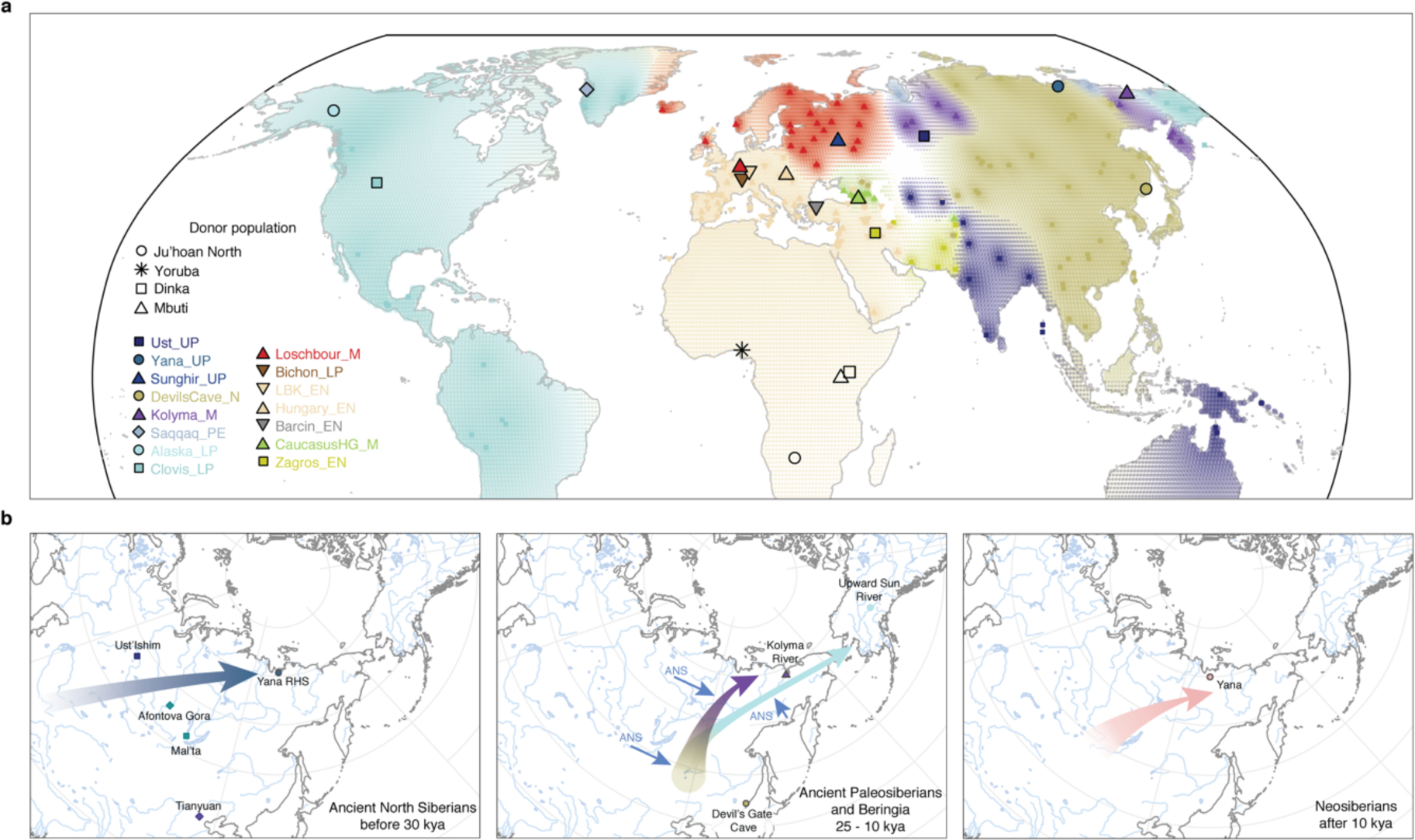
Genetic legacy of ancient Eurasians. **a,** World-wide map of top haplotype donations inferred by chromopainter. Each coloured symbol represents a modern recipient population, with the colour and shape indicating the donor population contributing the highest fraction of haplotypes to that recipient population. Geographic locations of donor populations used in this analysis (modern Africans and ancient Eurasians) are indicated by the corresponding larger symbols with black outline added. Extended regions of shared top donors are visualized by spatial interpolation of the respective donor population color. **b,**Major hypothesized migrations into northeast Siberia. Arrows indicate putative migrations giving rise to Ancient North Siberians (left), Ancient Paleosiberians (middle; possible ANS admixture scenarios indicated by small blue arrows) and Neosiberians (right). Key sample locations for the respective time slice are indicated with symbols.

Our Holocene transect reveals additional complexity in recent times, with evidence for further episodes of gene flow and localized population replacements. A striking example is found in the Lake Baikal region in southern Siberia, where the newly reported genomes from Ust’Belaya and recently published neighbouring Neolithic and Bronze Age sites show a succession of three distinct genetic ancestries over a ~6 ky time span. The earliest individuals show predominantly East Asian ancestry, closely related to the ancient individuals from DGC (Figure 1; Extended Data Fig. 6, 7). In the early Bronze Age (BA), we observe a resurgence of AP ancestry (up to ~50% ancestry fraction), as well as influence of West Eurasian Steppe ANE ancestry represented by the early BA individuals from Afanasievo in the Altai region (~10%) (Extended Data Fig. 7; Supplementary Data Table 3). This is consistent with previous reports of gene flow from an unknown ANE-related source into Lake Baikal hunter-gatherers^42^. Our results suggest a southward expansion of AP as a possible source, which is also consistent with the replacement of Y chromosome lineages observed at Lake Baikal, from predominantly haplogroup N in the Neolithic to haplogroup Q in the BA^42^. Finally, the most recent individual from Ust’Belaya, dated to ~600 years ago, falls along the Neosiberian cline, similar to the ~760 year-old ‘Young Yana’ individual from northeastern Siberia, demonstrating the widespread distribution of Neosiberian ancestry in the most recent epoch. We show that most populations on this cline can be modelled as predominantly East Asian, with varying proportions of West Eurasian ANE ancestry, with the largest proportions observed in later ancient and modern Altaian populations (Extended Data Fig. 7; Supplementary Data Table 3). Together, these findings show that by Holocene times there was considerable population movement and admixture throughout southern and eastern Siberia, with groups dispersing in multiple directions yet without clear evidence of the wholesale population replacement seen in earlier Pleistocene times.

We further investigated whether these processes of population flux were a more widespread phenomenon across all of Northern Eurasia. The striking spatial pattern of AP and East Asian ancestry in modern populations observed using chromosome painting (Fig. 4) strongly suggests that AP ancestry was once widespread, likely as far west as the Urals. At the western edge of northern Eurasia, genetic and strontium isotope data from ancient individuals at the Levänluhta site (Supplementary Information 1) documents the presence of Saami ancestry in Southern Finland in the Late Holocene 1.5 kya. This ancestry component is currently limited to the northern fringes of the region, mirroring the pattern observed for AP ancestry in northeastern Siberia. However, while the ancient Saami individuals harbour East Asian ancestry, we find that this is better modelled by DGC rather than AP, suggesting that AP influence was likely restricted to the eastern side of the Urals (Extended Data Fig. 7; Supplementary Data Table 3). Comparison of ancient Finns and Saami with their present-day counterparts reveals additional gene flow over the past 1.6 kya, with evidence for West Eurasian admixture into modern Saami. The ancient Finn from Levänluhta shows lower Siberian ancestry than modern Finns (Extended Data Table 2), therefore likely representing the Scandinavian component present in the dual-origin (Uralic/Scandinavian) gene pool of Finns.

## Discussion

Our findings reveal that the population history of northeastern Siberia is far more complex than has been inferred from the contemporary genetic record. It involved at a minimum three major population migrations and subsequent replacements, starting in the Late Pleistocene, with considerable smaller scale population fluxes throughout the Holocene. These three major changes are also clearly documented in the archaeological record. The initial movement into the region was by a now-extinct ANS population diversifying ~38 kya, soon after the basal West Eurasian and East Asian split, represented by the archaeological culture found at Yana RHS^4,53^. This finding is consistent with other studies that have shown this was a time of rapid expansion of early modern humans across Eurasia^44^. The arrival of peoples carrying ancestry from East Asia, and their admixture after the LGM with descendants of the ANS lineage ~20-18 kya, led to the replacement of the Yana population and the rise of the AP and Native American lineages. In the archaeological record this is demonstrated by the spread of microblade technology that accompanies the shrinkage of the mammoth habitat area in a northerly direction^10^. This group was in turn largely replaced by Neosiberians in the early and mid-Holocene. Our data suggest the Neosiberians received ANS ancestry indirectly through Kolyma ~11 kya, and possibly later gene flow from Bronze Age groups from the central Asian steppe (Afanasievo) after ~5 kya. Intriguingly, a signal of Australasian ancestry that has been observed in some Amazonian groups^24,28^ is not evident in any of the ancient Siberian or Beringian samples sequenced here, or in previous studies^41^.

We find that the first inhabitants of northeastern Siberia were not the ancestors of either Native Americans or modern Siberians. Instead, they were the earliest descendants of a deeply diverged early Eurasian lineage that apart from Yana is currently only represented by two ancient individuals from central Siberia (Mal’ta and Afontova Gora), widely separated from northeastern Siberia. The finding that these early groups were so highly distinct, and diverged so soon after 38 kya, suggests that the combined ANS/ANE lineages are much more complex than previously supposed. Although represented by just a few individuals, and lacking direct descendants, traces of their presence are recorded in ancient and modern genomes across northern Eurasia and the Americas. One of these groups, the “Ancient Paleosiberians”, likely also once had a wide geographic distribution across northern Eurasia. Its genetic legacy among present-day Siberians is more limited, restricted to groups in northeastern Siberia but also the Americas. Nonetheless, this distribution implies that the majority of Native American genetic ancestry likely originated in northeastern Siberia, rather than south-central Siberia, as inferred from modern mitochondrial and Y chromosome DNA^54^. The Neosiberians occupying much of the range previously inhabited by ANS and AP, represent a more recent arrival that originated further south. The replacement processes we have revealed for the northeastern portion of Siberia are mirrored in far western Eurasia by the regional replacement of the Saami people in the Late Holocene, suggesting similar processes likely took place in many other parts of the northern hemisphere.

## Methods

### Sample processing and DNA sequencing

The ancient DNA (aDNA) work was conducted in dedicated aDNA clean-room facilities at Centre for GeoGenetics, Natural History Museum, University of Copenhagen according to strict aDNA standards. DNA was extracted from the samples following established protocols^27,55^. Sequencing libraries were built from the extracts and amplified as previously described^56,57^ and sequenced on the Illumina platform. Raw reads were trimmed for Illumina adaptor sequences using AdapterRemoval^58^, and mapped to the human reference genome build 37 using BWA^59^ with seeding disabled^60^. Final analysis BAM files were obtained by discarding reads with mapping quality ≤ 30, removing PCR duplicates with MarkDuplicates (http://picard.sourceforge.net) and local realignment using GATK^61^.

### Authentication, mitochondrial DNA and chromosome Y analyses

Authentication for ancient DNA was carried out by examining fragment length distributions and nucleotide substitution patterns characteristic for ancient DNA damage using mapDamage^62^. Levels of contamination were estimated for all individuals on mitochondrial DNA sequences using schmutzi^63^, as well as on chromosome X for male individuals using angsd^64^. Mitochondrial DNA sequences were reconstructed using endoCaller from schmutzi^63^, and haplogroups assigned with HaploGrep^65^. Y chromosome haplogroups were assigned from reads overlapping SNPs included in the Y-DNA haplogroup tree from the International Society of Genetic Genealogy (ISOGG; http://www.isogg.org, version 13.37), as previously described^39^. Phylogenetic analysis was carried out on haploid SNP calls from high coverage individuals obtained with samtools/bcftools^66^, using RAxML^67^ with the ASC_GTRGAMMA model^44^.

### Analysis panels

Autosomal analyses were carried out on three analysis panels of ancient and modern individuals and different sets of SNPs. Panel 1 (“HO 1240K”) includes modern individuals from world-wide populations genotyped using the Affymetrix HumanOrigins array^14^, merged with ancient individuals with data from shotgun sequencing or genomic capture (the 1240K panel^68^). Panel 2 (“SGDP/CGG 2240K”) includes shotgun sequencing data for modern and ancient individuals, as well as selected ancient individuals with genomic capture, all genotyped at SNPs included in the 2240K capture panel^29,32^. Panel 3 (“CGG WGS”) includes all genome-wide SNPs genotyped across high coverage modern and ancient individuals with shotgun sequencing data. Genotyping was carried separately for each diploid individual using samtools/bcftools^66^, and filtered as previously described^39^ (Supplementary Information section 5). Pseudo-haploid genotypes for low-coverage ancient individuals were obtained by sampling a random high-quality read at each covered SNP position of the respective panels.

### Population structure and admixture modelling

Population structure was investigated with PCA using smartpca^69^. Principal components were inferred using modern as well as high coverage ancient individuals, followed by projection of low-coverage individuals using ‘*lsqproject’*. Genetic affinities of ancient and modern individuals were investigated with the *f*-statistic framework^70^, using ‘outgroup *f*_3_’ statistics for estimation of shared genetic drift^15^ as well as *f*_4_ statistics for allele sharing analyses. Admixture graph modelling was carried out using qpGraph, and outgroup-based estimation of admixture components using qpAdm from the ADMIXTOOLS package^70^ (Supplementary Information sections 7-9).

### Relatedness and identity-by-descent analyses

Relatedness among the ancient individuals was quantified using the kinship coefficient estimator implemented in KING^71^, obtained from a pairwise identity-by-state (IBS) matrix inferred with realSFS implemented in angsd^64^. Genomic segments homozygous-by-descent (HBD) and identical-by-descent (IBD) were inferred for all high-coverage individuals using IBDseq^72^ (Supplementary Information section 6).

### Demographic modelling

The parameters of alternative demographic scenarios were inferred based on the joint site frequency spectrum (SFS) by approximating the likelihood of a given model with coalescent simulations, using the composite likelihood method implemented in the fastsimcoal2 software^73^. Demographic modelling was carried out on selected ancient individuals from the “CGG WGS” panel, merged with a set of genomes of present-day individuals from the Simon’s Genome Diversity Project^37^. We discarded singleton SNPs for this analysis to minimize the influence of possible sequencing errors in the ancient individuals. Confidence intervals were obtained using a block-bootstrap approach, resampling blocks of 1Mb. Parameters in coalescent time were scaled to time in years assuming a mutation rate of 1.25 x 10^-8^ / generation / site^74^ and a generation time of 29 years^75^ (Supplementary Information section 10).

### Haplotype sharing analyses

Haplotype-based analyses of population structure were carried out using chromopainter^76^ on all individuals with diploid genotypes in both the “HO 1240K” and “WGS” datasets. We used shapeit^77^ to reconstruct phased haplotypes for each individual. Chromosome painting was then carried out as previously described^78^. We first estimated the parameters *N*_*e*_ and θ on a subset of individuals (chosen from diverse modern and ancient groups) and chromosomes (2, 9, 16, 22) using 10 iterations of the Expectation-Maximization (E-M) algorithm, separately for each dataset. Chromosome painting for inferring global population structure related to the ancient individuals was then performed by painting all non-African modern individuals as recipients, using African as well as high coverage ancient individuals as possible donors. Population structure was investigated by multidimensional scaling on the co-ancestry matrix obtained from chromopainter, both for length and number of shared chunks. For the analysis of the Siberian ancestry in present-day Athabascan groups a second analysis was carried out, by painting all Native American groups using modern Africans and ancient individuals from outside the Americas as potential donors. We quantified differential sharing of pairs of Native American populations A and B with a particular donor group using the symmetry statistic^24^

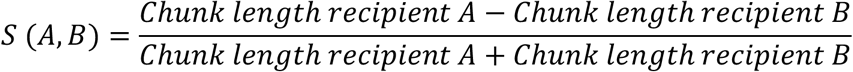

Standard errors were estimated using a block jacknife, dropping each of the 22 chromosomes in turn.

### Paleoclimate modelling

We used paleoclimatic modelling to identify regions with the most suitable climatic conditions, in steps of 1,000 years from 48 to 12kya. We collated a geo-referenced database of modern human fossil and archaeological dated remains, including 936 modern human occurrences across all time intervals. All paleoclimatic data were gridded to a 1×1 degree resolution, and all occurrences within a grid cell were aggregated to a single occurrence. Paleoclimatic conditions were simulated under the HadCM3 (Hadley Centre Coupled Model, version 3) Atmospheric– Ocean General Circulation Model (AOGCM), and we selected the three seasonal variables that maximized the climatic signal information: Autumn total precipitation, Summer average temperature and Autumn average temperature. An ensemble of seven different algorithms was used to characterise the climatic niche of modern humans, using the package “biomod2”. We validated the accuracy of the climatic suitability predictions using cross-validation within each time periods. To identify what regions had the most suitable climatic conditions across all time periods, from 48 to 12ka, we estimated the median suitability, and standard deviation, across time intervals for each grid cell (Supplementary Information section 12).

### Data availability

Sequence data were deposited in the European Nucleotide Archive (ENA) under accession: XXX

## Acknowledgements

This work was supported by The Lundbeck Foundation, The Danish National Research Foundation, and KU2016 (GeoGenetics). A. Y. F. was funded by the Russian Science Foundation (project No.14-50-00036). I.D. and V.S. were supported by a Swiss NSF grants 310030B-166605 and 31003A-143393 to L.E., and V.S. was further supported by Portuguese FCT (UID/BIA/00329/2013). V.P., E.Y.P., and P.A.N. are supported by Russian Science Foundation project N 16-18-10265-RNF. D.J.M. is supported by the Quest Archaeological Research Program.

---

**Extended Data Figure 1.**
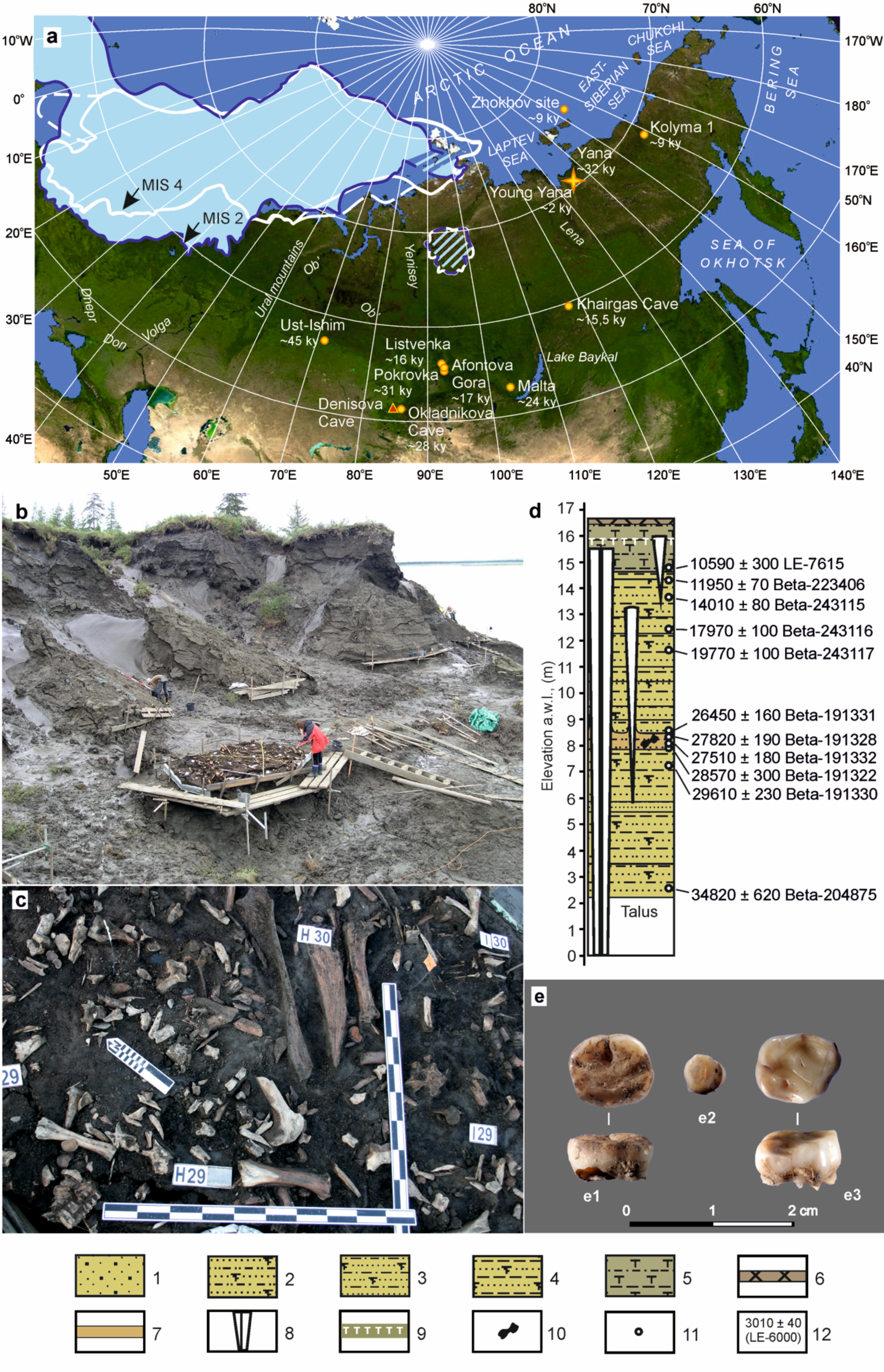
Geographical, chronological and archaeological context for the earliest human remains discovered in Northern Siberia. **a,** map of known 14C dated anatomically modern human fossils of late Pleistocene and early Holocene age (yellow dots) found in Siberia (Akimova et al. 2010; Alexeev 1998; Chikisheva et al. 2016; Fu et al. 2014; Khaldeyeva et al. 2016; Pitulko et al. 2015; Zubova and Chikisheva 2015) and Yana RHS finds (yellow star), Denisova Cave that yielded Neanderthal/Denisovan remains, red triangle (Chikisheva, Shunkov 2017; Reich et al. 2010) and the reconstructed maximum ice sheet extent at about 60,000 years ago (white line) and during the Last Glacial Maximum (LGM) around 20,000 years ago (ice-blue filling) (Hubberten et a. 2004; Svendsen et al. 2004); potentially glaciated areas are cross-hatched; **b,** general view of the Northern Point excavation area at the Yana site (Pitulko et al. 2004); **c,** cultural layer in H29 unit where the human tooth was found; d, cryolithological profile for Northern Point of Yana RSH (Pitulko et al. 2013); **e,** human teeth found during the excavations in unit 2V26, occlusal and lateral view (e1), unit X26 (e2), occlusal view, and H29 (e3), occlusal and lateral view, samples e2 (Yana 2 genome) and e3 (Yana 1 with high coverage (25.6X) genome sequence) are being used in this study. Legend for (c): 1 – sand with small pebbles; 2 – sandy silt; 3 – claey-sand silt; 4 – sandy-clayey silt; 5 – interbedding of clayey silt bands and sandy-clayey silt with beds and lenses of peat; 6 – soil-vegetable layer; 7 – culture layer; 8 – polygonal ice wedges; 9 – boundary of seasonal active layer; 10 – location of bones of Pleistocene animals sampled for 14С dating; 11 – location of 14С samples of plant remains; 12 – radiocarbon date and lab code.

**Extended Data Figure 2.**
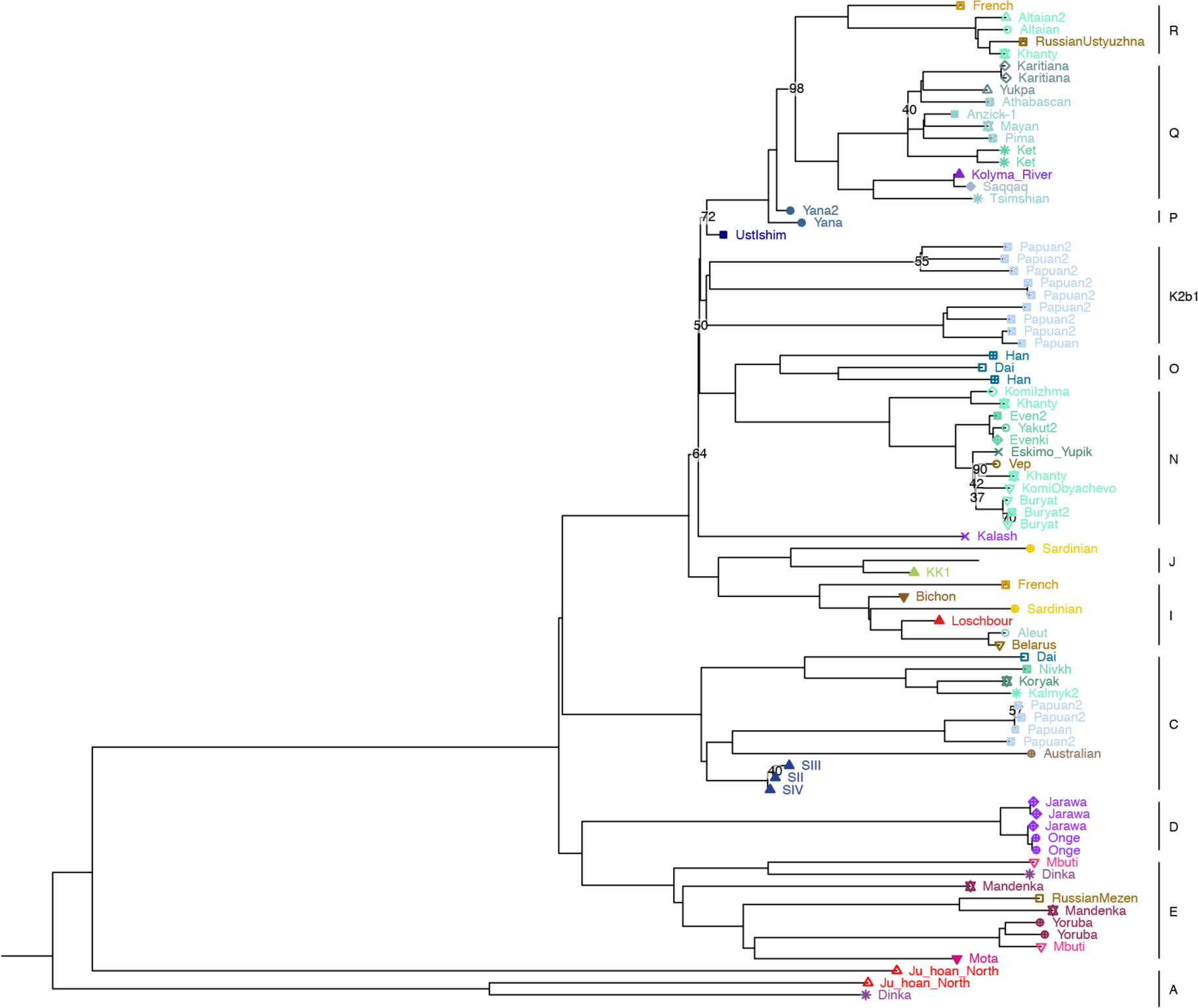
Y chromosome phylogeny for ancient and modern individuals, with major haplogroups highlighted

**Extended Data Figure 3.**
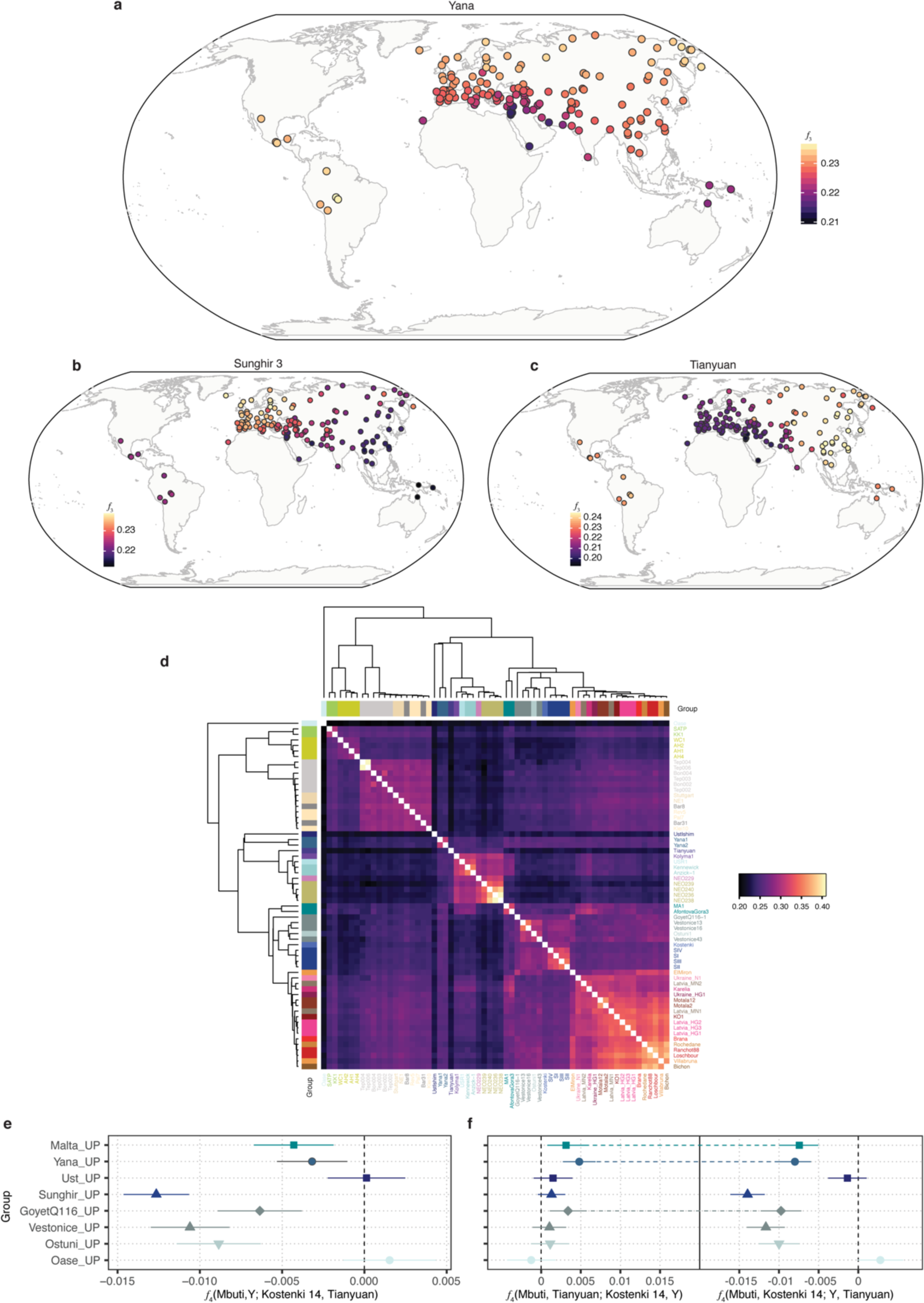
Genetic affinities of Yana. **a-c** Geographic heat maps depicting outgroup-*f*_3_ statistic for **a**, Yana1, **b**, Tianyuan and **c** Sunghir3 with world-wide populations. **d**, Heatmap showing genetic clustering of ancient individuals, based on pairwise outgroup-*f*_3_ statistic **e**, *f*_4_-statistics showing higher shared affinity between Yana and other selected UP groups to EWE (Kostenki). **f**, *f*_4_-statistics highlighting groups with affinities to both EEA and EWE (joined with dashed lines).

**Extended Data Figure 4.**
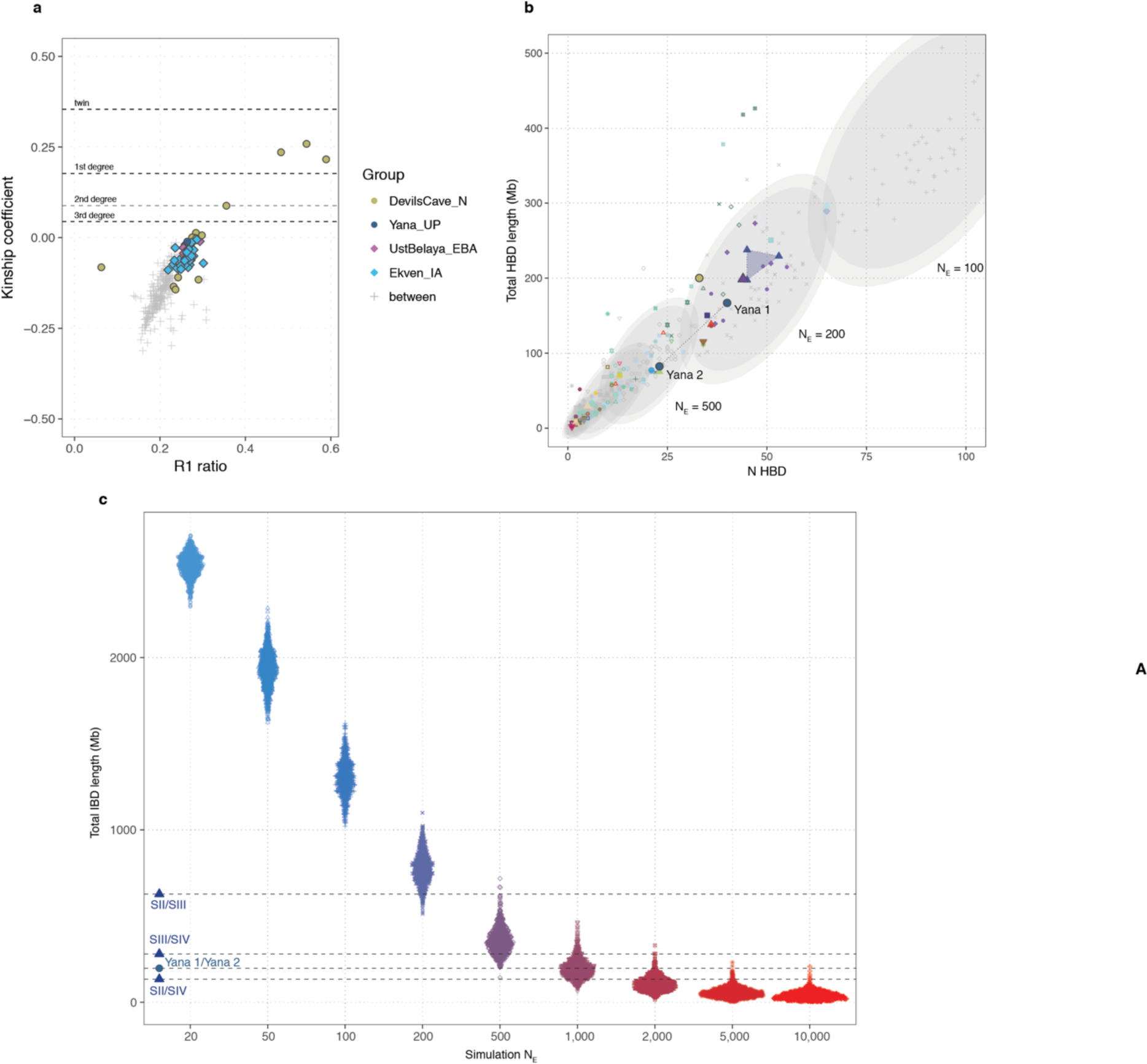
Relatedness and IBD. **a**, Kinship coefficients and R1 ratio (number of double heterozygous (Aa/Aa) sites divided by the total number of discordant genotypes) for newly reported ancient groups with multiple individuals per site. **b,**Number and length of HBD segments in ancient and modern individuals. Grey ellipses indicate confidence intervals for simulations of indicated effective population size. **c**, Distribution of total IBD lengths for simulations of varying effective population sizes. Observed values for pairs from Sunghir and Yana are indicated by dashed lines.

**Extended Data Figure 5.**
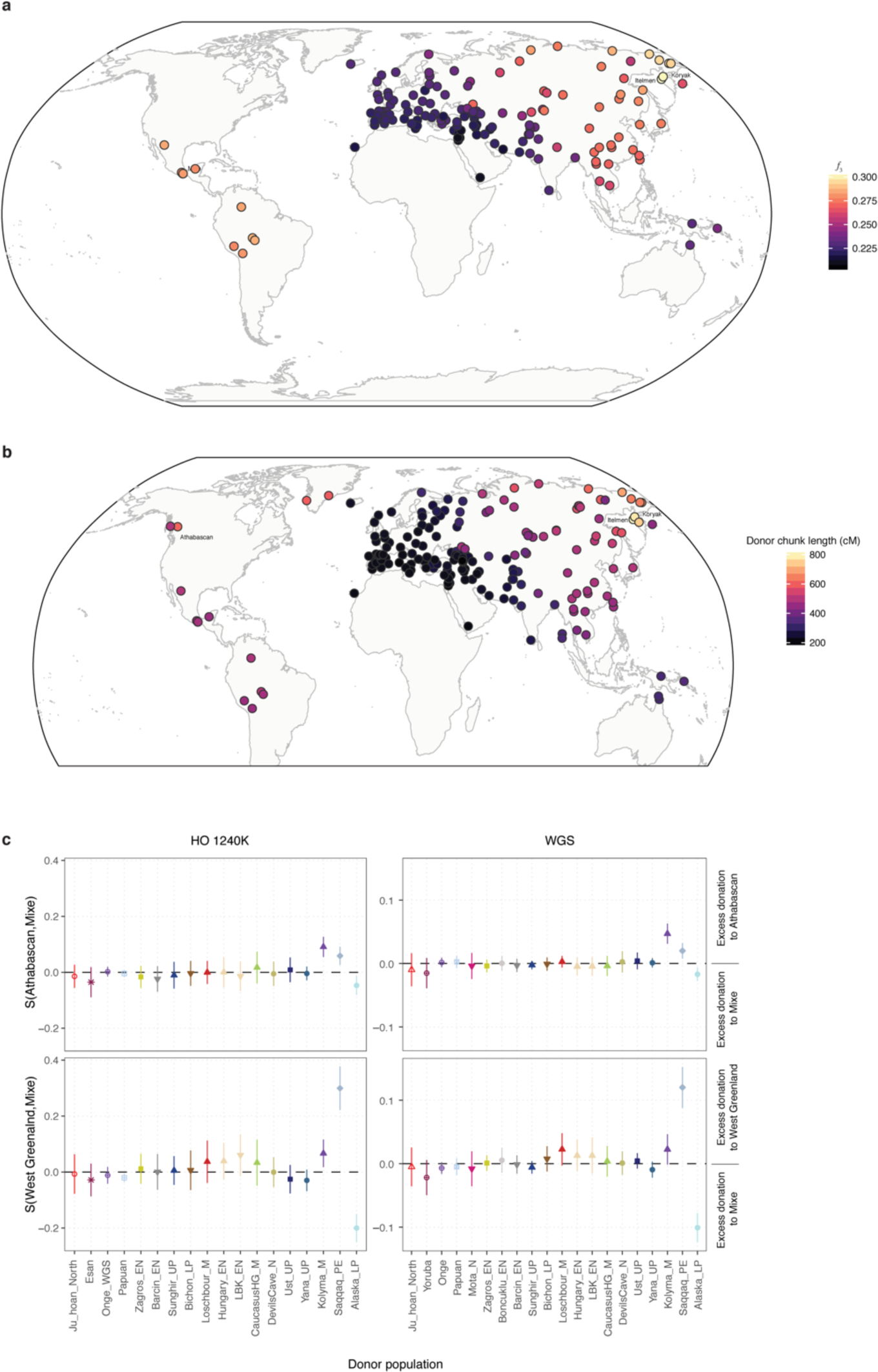
Genetic affinities of Kolyma1. **a, b** Geographic heat maps depicting genetic affinities of Kolyma individual with world-wide populations using **(a)** outgroup-*f*_3_ statistics and **(b)** total length of haplotype chunks donated by modern populations in chromosome painting. **c**, chromosome painting symmetry statistic contrasting the total length of haplotypes donated from ancient and modern non-American donor groups to pairs of American populations. The top panels show greater excess in donations to Athabascans from Kolyma1. The bottom panel shows the same statistic for West Greenland Inuit, a population with known affinity to Paleoeskimos, reflected in the excess donations observed from Saqqaq.

**Extended Data Figure 6.**
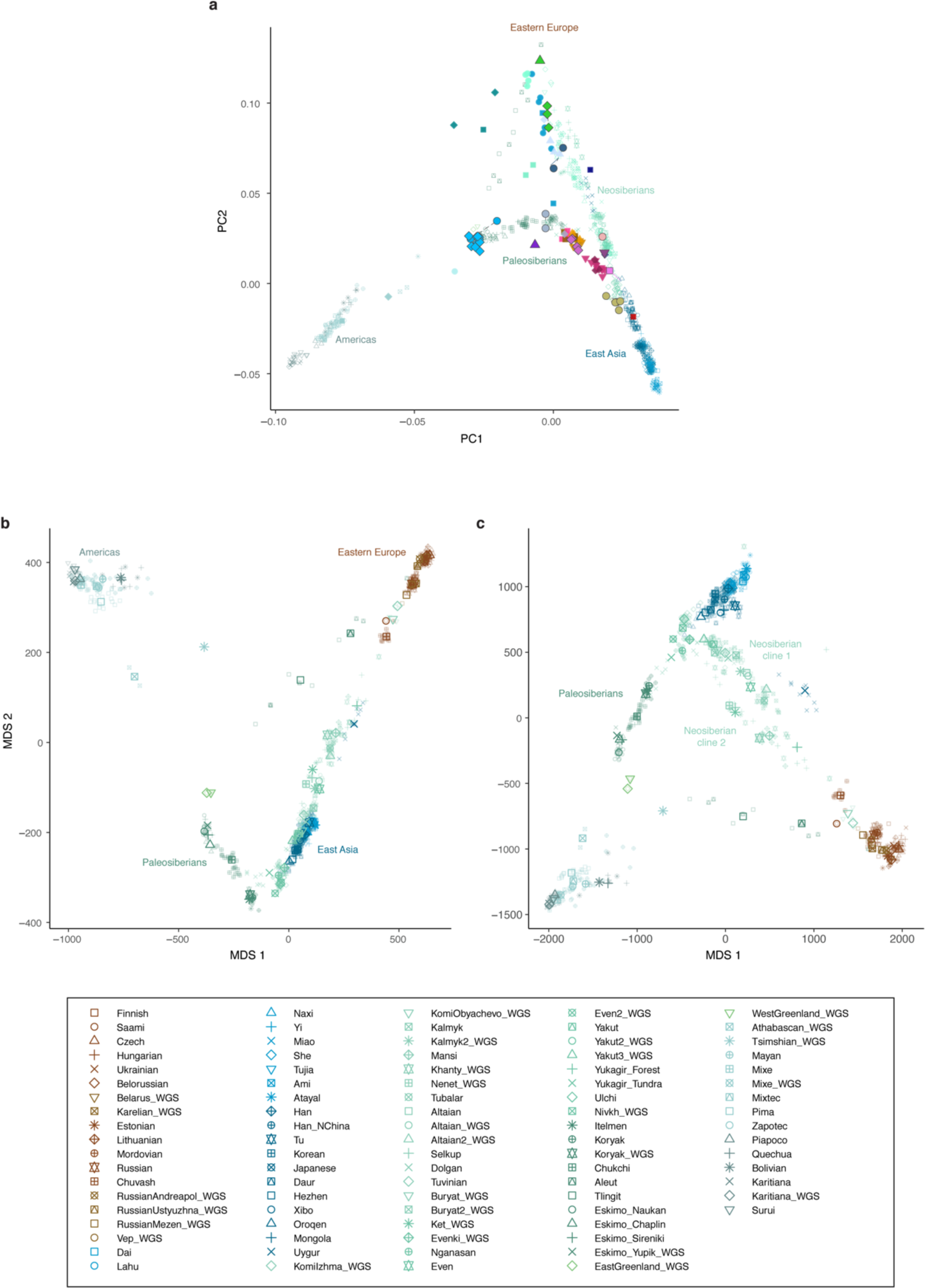
Genetic diversity in Northern Eurasia related to ancient genomes. **a**, PCA of modern individuals with ancient groups projected. **b, c** MDS plots of modern populations, obtained from the chromosome painting co-ancestry matrix using modern Africans and high coverage ancient individuals as donors. **(b)** total length of chunks. **(c)** total number of chunks

**Extended Data Figure 7.**
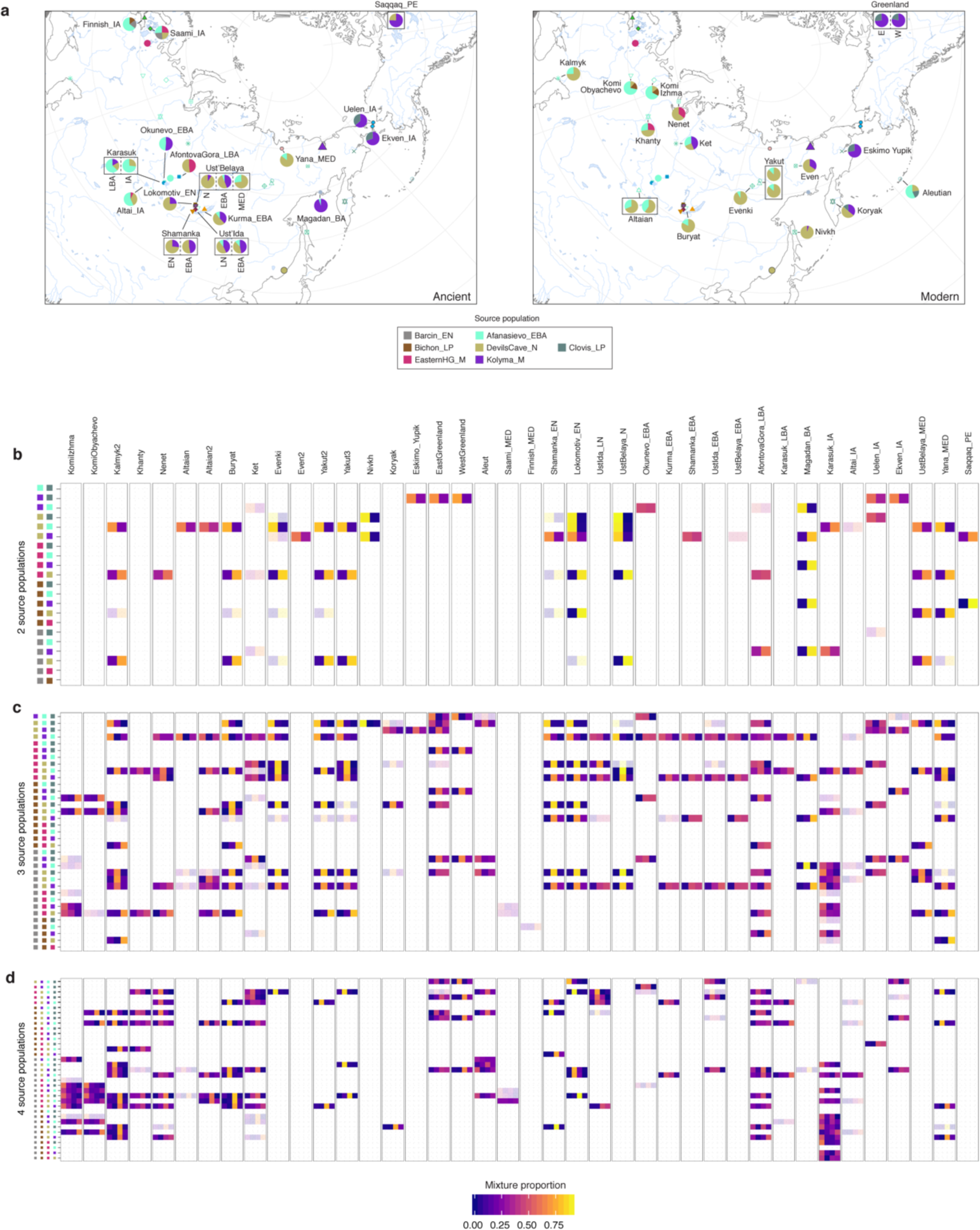
Admixture modelling using qpAdm. **a**, Maps showing locations and ancestry proportions of ancient (left) and modern (right) groups. **b-d**, Ancestry proportions and fit for all possible 2-way (b), 3-way (c) and 4-way (d) reference population combinations. Transparency shading indicates model fit, with lighter transparency indicating models accepted with 0.05 > p ≥ 0.01.

**Extended Data Figure 8.**
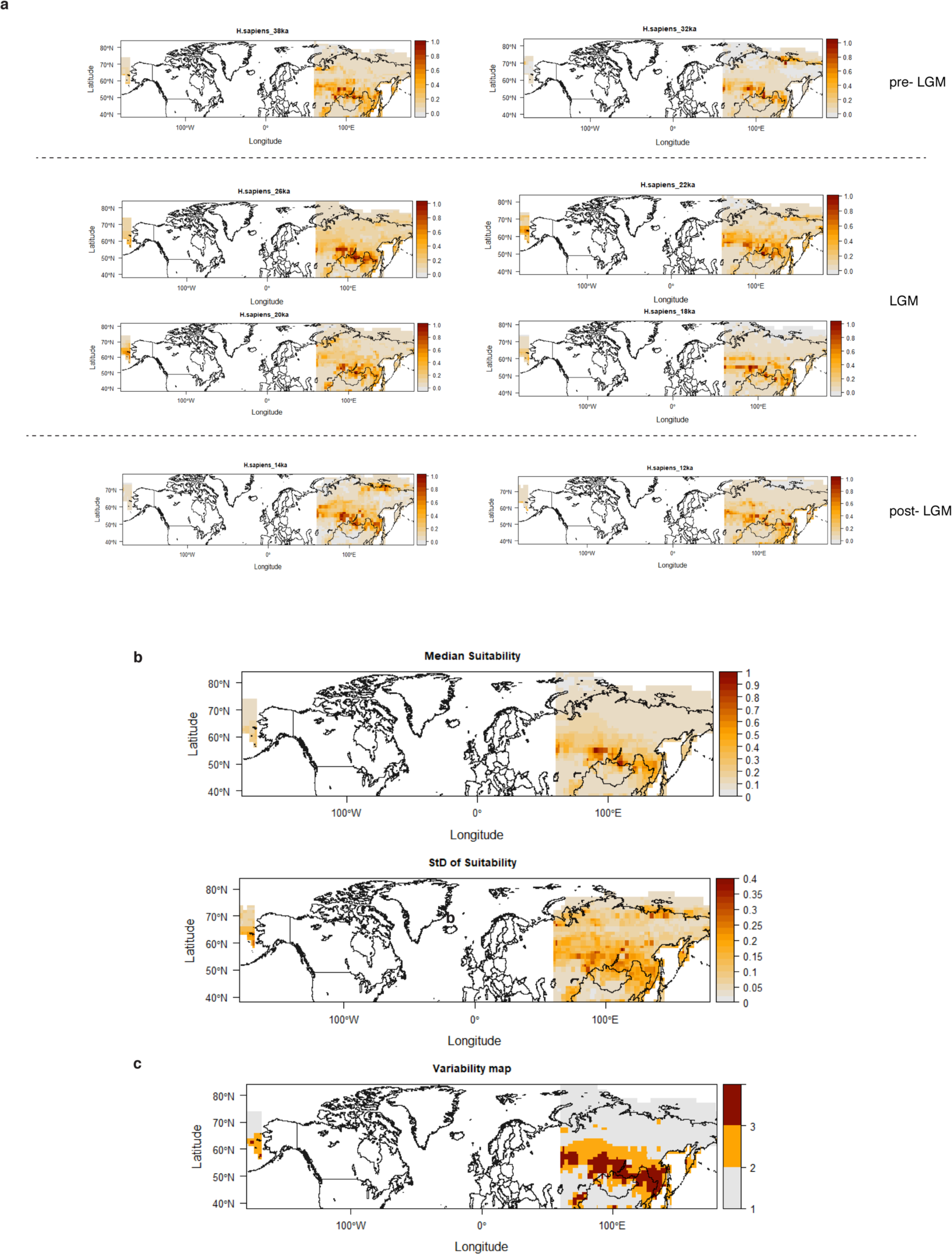
Paleo-climatic niche modelling. Maps showing climatically suitable regions for human occupation across temporal and spatial dimensions. Projections are bounded between 60 E to 180 E and from 38 N to 80N. Colour-key represents suitability values, with darker (lighter) colours corresponding to higher (lower) suitability values. **a**, Examples of climatic suitability for human occupation for different time slices. **b,**Median and standard deviation of climatic suitability across 23 climatic periods of millennial or bimillennial time resolution. **c**, Regions highly climatically suitable for humans (red), low (grey), and regions with both periods of high and low suitability (orange)

**Extended Data Table 1.**
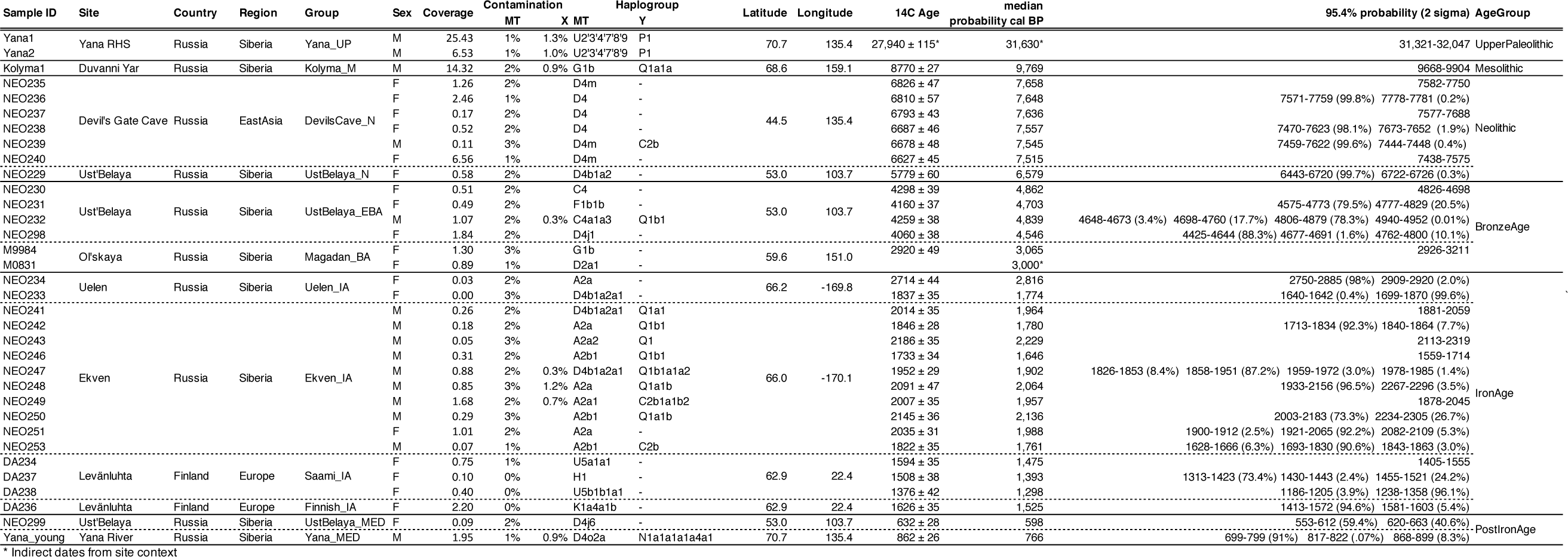
Newly reported ancient individuals

**Extended Data Table 2.**
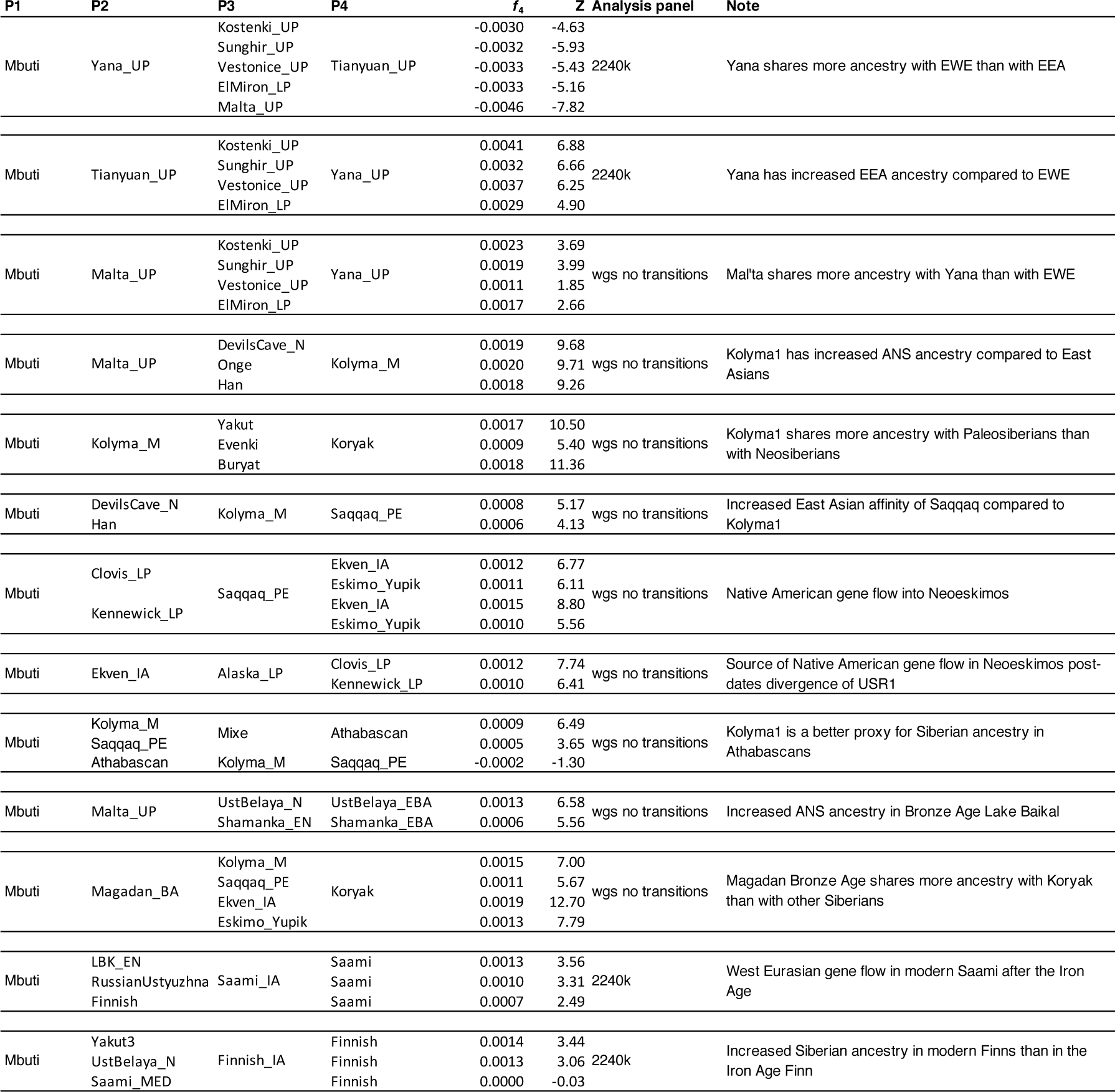
Key f-statistics.

## References

1. Fedorova, S. A. et al. Autosomal and uniparental portraits of the native populations of Sakha (Yakutia): implications for the peopling of Northeast Eurasia. BMC Evolutionary Biology 13, 127 (2013).

2. Pugach, I. et al. The Complex Admixture History and Recent Southern Origins of Siberian Populations. Mol Biol Evol msw055 (2016). doi:10.1093/molbev/msw055

3. Wong, E. H. M. et al. Reconstructing genetic history of Siberian and Northeastern European populations. Genome Res. 27, 1–14 (2017).

4. Pitulko, V. V. et al. The Yana RHS Site: Humans in the Arctic Before the Last Glacial Maximum. Science 303, 52–56 (2004).

5. Pitulko, V. V., Nikolskiy, P. A., Basilyan, A. & Pavlova, E. Y. Human Habitation in Arctic Western Beringia Prior to the LGM. in Paleoamerican Odyssey (eds. Graf, K. E., Ketron, C. V. & Waters, M. R.) (Texas A&M University Press, 2014).

6. Pitulko, V. V. et al. Early human presence in the Arctic: Evidence from 45,000-year-old mammoth remains. Science 351, 260–263 (2016).

7. Pitulko, V., Pavlova, E. & Nikolskiy, P. Revising the archaeological record of the Upper Pleistocene Arctic Siberia: Human dispersal and adaptations in MIS 3 and 2. Quaternary Science Reviews 165, 127–148 (2017).

8. Rasmussen, S. O. et al. A stratigraphic framework for abrupt climatic changes during the Last Glacial period based on three synchronized Greenland ice-core records: refining and extending the INTIMATE event stratigraphy. Quaternary Science Reviews 106, 14–28 (2014).

9. Derevîanko, A. P., Powers, W. R. & Shimkin, D. B. The Paleolithic of Siberia: New Discoveries and Interpretations. (Institute of Archaeology and Ethnography, Siberian Division, Russian Academy of Sciences, 1998).

10. Pitulko, V. V. & Nikolskiy, P. A. The extinction of the woolly mammoth and the archaeological record in Northeastern Asia. World Archaeology 44, 21–42 (2012).

11. Siska, V. et al. Genome-wide data from two early Neolithic East Asian individuals dating to 7700 years ago. Science Advances 3, e1601877 (2017).

12. Rasmussen, M. et al. Ancient human genome sequence of an extinct Palaeo-Eskimo. Nature 463, 757–762 (2010).

13. Meyer, M. et al. A High-Coverage Genome Sequence from an Archaic Denisovan Individual. Science 338, 222–226 (2012).

14. Lazaridis, I. et al. Ancient human genomes suggest three ancestral populations for present-day Europeans. Nature 513, 409–413 (2014).

15. Raghavan, M. et al. Upper Palaeolithic Siberian genome reveals dual ancestry of Native Americans. Nature 505, 87–91 (2014).

16. Rasmussen, M. et al. The genome of a Late Pleistocene human from a Clovis burial site in western Montana. Nature 506, 225–229 (2014).

17. Raghavan, M. et al. The genetic prehistory of the New World Arctic. Science 345, 1255832 (2014).

18. Prüfer, K. et al. The complete genome sequence of a Neanderthal from the Altai Mountains. Nature 505, 43–49 (2014).

19. Fu, Q. et al. Genome sequence of a 45,000-year-old modern human from western Siberia. Nature 514, 445–449 (2014).

20. Seguin-Orlando, A. et al. Genomic structure in Europeans dating back at least 36,200 years. Science 346, 1113–1118 (2014).

21. Olalde, I. et al. Derived immune and ancestral pigmentation alleles in a 7,000-year-old Mesolithic European. Nature 507, 225–228 (2014).

22. Gamba, C. et al. Genome flux and stasis in a five millennium transect of European prehistory. Nat Commun 5, 5257 (2014).

23. Rasmussen, M. et al. The ancestry and affiliations of Kennewick Man. Nature 523, 455–458 (2015).

24. Skoglund, P. et al. Genetic evidence for two founding populations of the Americas. Nature 525, 104–108 (2015).

25. Llorente, M. G. et al. Ancient Ethiopian genome reveals extensive Eurasian admixture throughout the African continent. Science 350, 820–822 (2015).

26. Ayub, Q. et al. The Kalash Genetic Isolate: Ancient Divergence, Drift, and Selection. The American Journal of Human Genetics 96, 775–783 (2015).

27. Allentoft, M. E. et al. Population genomics of Bronze Age Eurasia. Nature 522, 167–172 (2015).

28. Raghavan, M. et al. Genomic evidence for the Pleistocene and recent population history of Native Americans. Science 349, aab3884 (2015).

29. Fu, Q. et al. An early modern human from Romania with a recent Neanderthal ancestor. Nature 524, 216–219 (2015).

30. Jones, E. R. et al. Upper Palaeolithic genomes reveal deep roots of modern Eurasians. Nat Commun 6, 8912 (2015).

31. Broushaki, F. et al. Early Neolithic genomes from the eastern Fertile Crescent. Science 353, 499–503 (2016).

32. Fu, Q. et al. The genetic history of Ice Age Europe. Nature 534, 200–205 (2016).

33. Mondal, M. et al. Genomic analysis of Andamanese provides insights into ancient human migration into Asia and adaptation. Nat Genet 48, 1066–1070 (2016).

34. Kılınç, G. M. et al. The Demographic Development of the First Farmers in Anatolia. Curr Biol 26, 2659–2666 (2016).

35. Jeong, C. et al. Long-term genetic stability and a high-altitude East Asian origin for the peoples of the high valleys of the Himalayan arc. PNAS 113, 7485–7490 (2016).

36. Hofmanová, Z. et al. Early farmers from across Europe directly descended from Neolithic Aegeans. PNAS 113, 6886–6891 (2016).

37. Mallick, S. et al. The Simons Genome Diversity Project: 300 genomes from 142 diverse populations. Nature 538, 201–206 (2016).

38. Jones, E. R. et al. The Neolithic Transition in the Baltic Was Not Driven by Admixture with Early European Farmers. Current Biology 27, 576–582 (2017).

39. Sikora, M. et al. Ancient genomes show social and reproductive behavior of early Upper Paleolithic foragers. Science 358, 659–662 (2017).

40. Yang, M. A. et al. 40,000-Year-Old Individual from Asia Provides Insight into Early Population Structure in Eurasia. Current Biology 27, 3202–3208.e9 (2017).

41. Moreno-Mayar, J. V. et al. Terminal Pleistocene Alaskan genome reveals first founding population of Native Americans. Nature 553, 203 (2018).

42. Damgaard, P. de B. et al. The first horse herders and the impact of early Bronze Age steppe expansions into Asia. Science eaar7711 (2018). doi:10.1126/science.aar7711

43. Dulik, M. C. et al. Y-chromosome analysis reveals genetic divergence and new founding native lineages in Athapaskan- and Eskimoan-speaking populations. PNAS 109, 8471–8476 (2012).

44. Poznik, G. D. et al. Punctuated bursts in human male demography inferred from 1,244 worldwide Y-chromosome sequences. Nature Genetics 48, 593 (2016).

45. Posth, C. et al. Pleistocene Mitochondrial Genomes Suggest a Single Major Dispersal of Non-Africans and a Late Glacial Population Turnover in Europe. Current Biology 26, 827–833 (2016).

46. Lipson, M. & Reich, D. A. Working Model of the Deep Relationships of Diverse Modern Human Genetic Lineages Outside of Africa. Mol Biol Evol 34, 889–902 (2017).

47. Haak, W. et al. Massive migration from the steppe was a source for Indo-European languages in Europe. Nature 522, 207–211 (2015).

48. Hoffecker, J. F., Elias, S. A., O’Rourke, D. H., Scott, G. R. & Bigelow, N. H. Beringia and the global dispersal of modern humans. Evolutionary Anthropology: Issues, News, and Reviews 25, 64–78 (2016).

49. Reich, D. et al. Reconstructing Native American population history. Nature 488, 370–374 (2012).

50. Loh, P.-R. et al. Inferring Admixture Histories of Human Populations Using Linkage Disequilibrium. Genetics 193, 1233–1254 (2013).

51. Flegontov, P. et al. Genomic study of the Ket: a Paleo-Eskimo-related ethnic group with significant ancient North Eurasian ancestry. Scientific Reports 6, 20768 (2016).

52. Flegontov, P. et al. Paleo-Eskimo genetic legacy across North America. bioRxiv 203018 (2017). doi:10.1101/203018

53. Pitulko, V. V., Pavlova, E. Y., Nikolskiy, P. A. & Ivanova, V. V. The oldest art of the Eurasian Arctic: personal ornaments and symbolic objects from Yana RHS, Arctic Siberia. Antiquity 86, 642–659 (2012).

54. Dulik, M. C. et al. Mitochondrial DNA and Y Chromosome Variation Provides Evidence for a Recent Common Ancestry between Native Americans and Indigenous Altaians. The American Journal of Human Genetics 90, 229–246 (2012).

55. Damgaard, P. B. et al. Improving access to endogenous DNA in ancient bones and teeth. Sci Rep 5, 11184 (2015).

56. Orlando, L. et al. Recalibrating Equus evolution using the genome sequence of an early Middle Pleistocene horse. Nature 499, 74 (2013).

57. Dabney, J. & Meyer, M. Length and GC-biases during sequencing library amplification: a comparison of various polymerase-buffer systems with ancient and modern DNA sequencing libraries. BioTechniques 52, 87–94 (2012).

58. Schubert, M., Lindgreen, S. & Orlando, L. AdapterRemoval v2: rapid adapter trimming, identification, and read merging. BMC Res Notes 9, (2016).

59. Li, H. & Durbin, R. Fast and accurate short read alignment with Burrows–Wheeler transform. Bioinformatics 25, 1754–1760 (2009).

60. Schubert, M. et al. Improving ancient DNA read mapping against modern reference genomes. BMC Genomics 13, 178 (2012).

61. DePristo, M. A. et al. A framework for variation discovery and genotyping using next-generation DNA sequencing data. Nat Genet 43, 491–498 (2011).

62. Jónsson, H., Ginolhac, A., Schubert, M., Johnson, P. L. F. & Orlando, L. mapDamage2.0: fast approximate Bayesian estimates of ancient DNA damage parameters. Bioinformatics 29, 1682–1684 (2013).

63. Renaud, G., Slon, V., Duggan, A. T. & Kelso, J. Schmutzi: estimation of contamination and endogenous mitochondrial consensus calling for ancient DNA. Genome Biol 16, 224 (2015).

64. Korneliussen, T. S., Albrechtsen, A. & Nielsen, R. ANGSD: Analysis of Next Generation Sequencing Data. BMC Bioinformatics 15, 356 (2014).

65. Weissensteiner, H. et al. HaploGrep 2: mitochondrial haplogroup classification in the era of high-throughput sequencing. Nucleic Acids Res 44, W58–W63 (2016).

66. Li, H. A statistical framework for SNP calling, mutation discovery, association mapping and population genetical parameter estimation from sequencing data. Bioinformatics 27, 2987–2993 (2011).

67. Stamatakis, A. RAxML version 8: a tool for phylogenetic analysis and post-analysis of large phylogenies. Bioinformatics 30, 1312–1313 (2014).

68. Mathieson, I. et al. Genome-wide patterns of selection in 230 ancient Eurasians. Nature 528, 499–503 (2015).

69. Patterson, N., Price, A. L. & Reich, D. Population Structure and Eigenanalysis. PLoS Genet 2, e190 (2006).

70. Patterson, N. et al. Ancient Admixture in Human History. Genetics 192, 1065–1093 (2012).

71. Manichaikul, A. et al. Robust relationship inference in genome-wide association studies. Bioinformatics 26, 2867–2873 (2010).

72. Browning, B. L. & Browning, S. R. Detecting Identity by Descent and Estimating Genotype Error Rates in Sequence Data. Am J Hum Genet 93, 840–851 (2013).

73. Excoffier, L., Dupanloup, I., Huerta-Sánchez, E., Sousa, V. C. & Foll, M. Robust Demographic Inference from Genomic and SNP Data. PLoS Genet 9, e1003905 (2013).

74. Scally, A. The mutation rate in human evolution and demographic inference. Current Opinion in Genetics & Development 41, 36–43 (2016).

75. Fenner, J. N. Cross-cultural estimation of the human generation interval for use in genetics-based population divergence studies. Am. J. Phys. Anthropol. 128, 415–423 (2005).

76. Lawson, D. J., Hellenthal, G., Myers, S. & Falush, D. Inference of Population Structure using Dense Haplotype Data. PLoS Genet 8, e1002453 (2012).

77. Delaneau, O., Zagury, J.-F. & Marchini, J. Improved whole-chromosome phasing for disease and population genetic studies. Nat Meth 10, 5–6 (2013).

78. Hellenthal, G. et al. A Genetic Atlas of Human Admixture History. Science 343, 747–751 (2014).

79. Malaspinas, A.-S. et al. A genomic history of Aboriginal Australia. Nature 538, 207–214 (2016).

